# Deep learning enables the atomic structure determination of the Fanconi Anemia core complex from cryoEM

**DOI:** 10.1101/2020.05.01.072751

**Authors:** Daniel P. Farrell, Ivan Anishchenko, Shabih Shakeel, Anna Lauko, Lori A. Passmore, David Baker, Frank DiMaio

**Affiliations:** Department of Biochemistry, University of Washington, Seattle WA 98105, USA; Institute for Protein Design, University of Washington, Seattle WA 98105, USA; MRC Laboratory of Molecular Biology, Cambridge, UK

## Abstract

Cryo-electron microscopy of protein complexes often leads to moderate resolution maps (4-8 Å), with visible secondary structure elements but poorly resolved loops, making model-building challenging. In the absence of high-resolution structures of homologues, only coarse-grained structural features are typically inferred from these maps, and it is often impossible to assign specific regions of density to individual protein subunits. This paper describes a new method for overcoming these difficulties that integrates predicted residue distance distributions from a deep-learned convolutional neural network, computational protein folding using Rosetta, and automated EM-map-guided complex assembly. We apply this method to a 4.6 Å resolution cryoEM map of Fanconi Anemia core complex (FAcc), an E3 ubiquitin ligase required for DNA interstrand crosslink repair, which was previously challenging to interpret as it is comprised of 6557 residues, only 1897 of which are covered by homology models. In the published structure built from this map, only 387 residues could be assigned to specific subunits. By building and placing into density 42 deep-learning guided models containing 4795 residues not included in the previously published structure, we are able to determine an almost-complete atomic model of FAcc, in which 5182 of the 6557 residues were placed. The resulting model is consistent with previously published biochemical data, and facilitates interpretation of disease related mutational data. We anticipate that our approach will be broadly useful for cryoEM structure determination of large complexes containing many subunits for which there are no homologues of known structure.

## Introduction

With the advent of direct electron detectors and advances in image processing software there has been an influx of large protein complex structures determined with cryoelectron microscopy (cryoEM). These technologies allow the structural characterization of protein assemblies that have eluded X-ray crystallography, and have led to maps with resolutions that allow atomic models to be built directly (3.3-4.6 Å or better) (Y. Cheng & Walz, 2009; Hryc et al., 2011) or lower subnanometer resolutions (∼5-9 Å) that can be interpreted by fitting of existing models. CryoEM data are noisy and structure determination requires a large number of particle images to be averaged together. This averaging, when combined with complications such as image misclassification, highly heterogeneous samples, or a limited number of sample views, typically limits the resolutions that can be attained (Lyumkis, 2019). This makes map interpretation difficult, and has necessitated the development of a number of tools for model-building and refinement into such cryoEM maps (Bonomi et al., 2019; Chen et al., 2016; Segura et al., 2016; Terashi & Kihara, 2018; Terwilliger et al., 2018; van Zundert et al., 2015).

In the absence of homologous structural information, cryoEM structures at intermediate resolutions are often largely “uninterpretable;” that is, while secondary structures may be identified and domains can often be roughly segmented, atomic-level information may not be accurately inferred (Gatsogiannis et al., 2013; Janssen et al., 2015; Kube et al., 2014; Stuttfeld et al., 2018). At best, a combination of secondary structure placement and sequence-based secondary structure prediction can lead to low-resolution complete or partial backbone trace models (L. Cheng et al., 2010; Snijder et al., 2017). Furthermore, while computational tools exist for modelling in maps at these resolutions (Bonomi et al., 2019; Kovacs et al., 2018; Segura et al., 2016; van Zundert et al., 2015; Webb et al., 2018), no tool is capable of inferring such structures *de novo*. Finally, while co-evolution information can provide valuable structural information (Kim et al., 2014; Nugent & Jones, 2012; Ovchinnikov et al., 2014), the availability of large numbers of sequences makes the method of limited applicability, though it has been used in determination of some cryoEM structures (Klink et al., 2020; Y.-J. Park et al., 2018; Schoebel et al., 2017).

In this manuscript, we take advantage of recent advances in protein structure prediction which employ deep convolutional neural networks to predict protein contacts or pairwise distances from multiple sequence alignments (Kandathil et al., 2019; Nugent & Jones, 2012; Senior et al., 2020; Xu, 2019; Yang et al., 2020; Zheng et al., 2019). We combine predictions from *trRosetta (Yang et al., 2020)*, which uses a deep residual-convolutional neural network to predict both distance and orientation between all pairs of residues in a protein, and a fast model building protocol that utilizes the results from the network to constraint folding. We then dock models generated using this approach into cryo-EM maps. The experimental EM data and deep-learning based structure predictions are synergistic: the deep-learned predictions serve the same role as high-resolution structures of homologues, informing the topology of individual domains and making the search space manageable, while the EM data, addresses two weaknesses in contact-guided prediction: it validates the accuracy of contact-guided predictions, and secondly, it provides information on the quaternary structure of complexes.

We illustrate the effectiveness of this approach by building an atomistic model of the *Gallus gallus* (chicken) Fanconi Anemia core complex (FAcc), guided by a recently published heterogeneous 4.6 Å single-particle cryoEM reconstruction and cross-linking mass spectrometry data. In previous work, crystal structures of FANCF, FANCE and FANCL were docked, and secondary structural elements were placed into the map (Shakeel et al., 2019). In contrast, here we are able to generate an atomistic model for nearly all of the complex. This method overcomes limitations of direct interpretation of the cryoEM map, including a lack of recognizable homology to proteins of known structure for the majority of subunits, and the relatively low resolution of substantial portions of the complex. The novel structural information provided by trRosetta-predicted distance-distributions enables accurate topology-level predictions for domains and subunits with no recognizable homology. By combining these trRosetta predictions (and Rosetta density-guided modelling tools (Wang et al., 2016)) with subnanometer resolution cryoEM data, we are able to infer a nearly-complete FAcc model, providing key insights into the function and organization of this complex.

## Results

A full description of our methodology is described in below and in the *Methods*. Briefly, Figure 1 illustrates how we determine complex structures using trRosetta models. The protocol is a five-stage process where we first generate multiple sequence alignments (MSAs) for the target proteins and use these to manually segment sequences into domains. Second, trRosetta is used to build models corresponding to these domains. Third, using Rosetta’s *dock_into_density*, we search the cryoEM reconstruction for the best-matching placements of each domain model. In the fourth step, we take all docked results in addition to crosslinking-mass spectrometry (XL-MS) data, and using Monte Carlo sampling of domain assignments in density, we find the arrangement of (and choice of) domain models maximizing the agreement with the cryoEM and XL-MS data. Finally, using RosettaCM we rebuild the connections between domains, and refine the entire complex against the cryoEM map.

**Figure 1.**
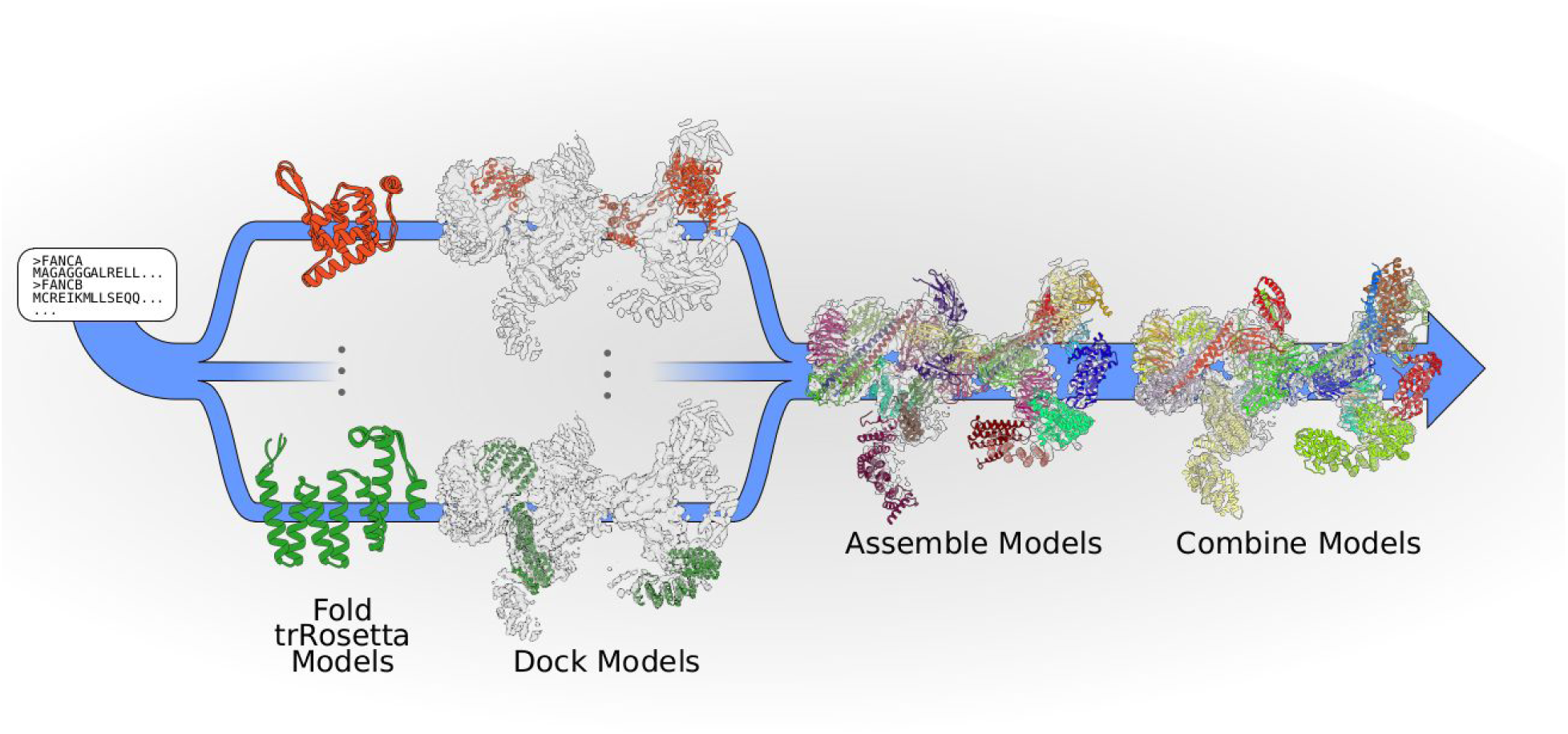
An overview of the workflow for modelling the FAcc. Initially, multiple sequence alignments (MSAs) of all protein sequences are generated, sequences are segmented into domains using the MSAs, and individual domains are folded using trRosetta. These domains are individually docked into the cryoEM density. Monte Carlo sampling finds the domain assignment maximally consistent with the density. Finally, linkers between domains are sampled and the entire structure is refined.

We illustrate the power of trRosetta predictions by applying this approach to build an atomic model into the recently determined cryoEM reconstruction of the Fanconi Anemia core complex (FAcc) (Shakeel et al., 2019). These data were obtained from a fully recombinant complex after overexpression of 8 protein subunits (FANCA, FANCB, FANCC, FANCE, FANCF, FANCG, FANCL, and FAAP100) in insect cells. A 3D reconstruction at an overall resolution of 4.6 Å and cross-linking mass spectrometry data were obtained. We previously identified secondary structure elements within the majority of the cryoEM map, and fit in homology models (Figure 2A). Using crosslinking, native and hydrogen-deuterium exchange mass spectrometry, as well as EM of purified subcomplexes we identified the general locations of all components except FANCA. However, in this previous work, we were only able to confidently determine residue assignments for FANCL. To gain further insight into the molecular mechanisms of FAcc, atomic models of all subunits are required.

**Figure 2.**
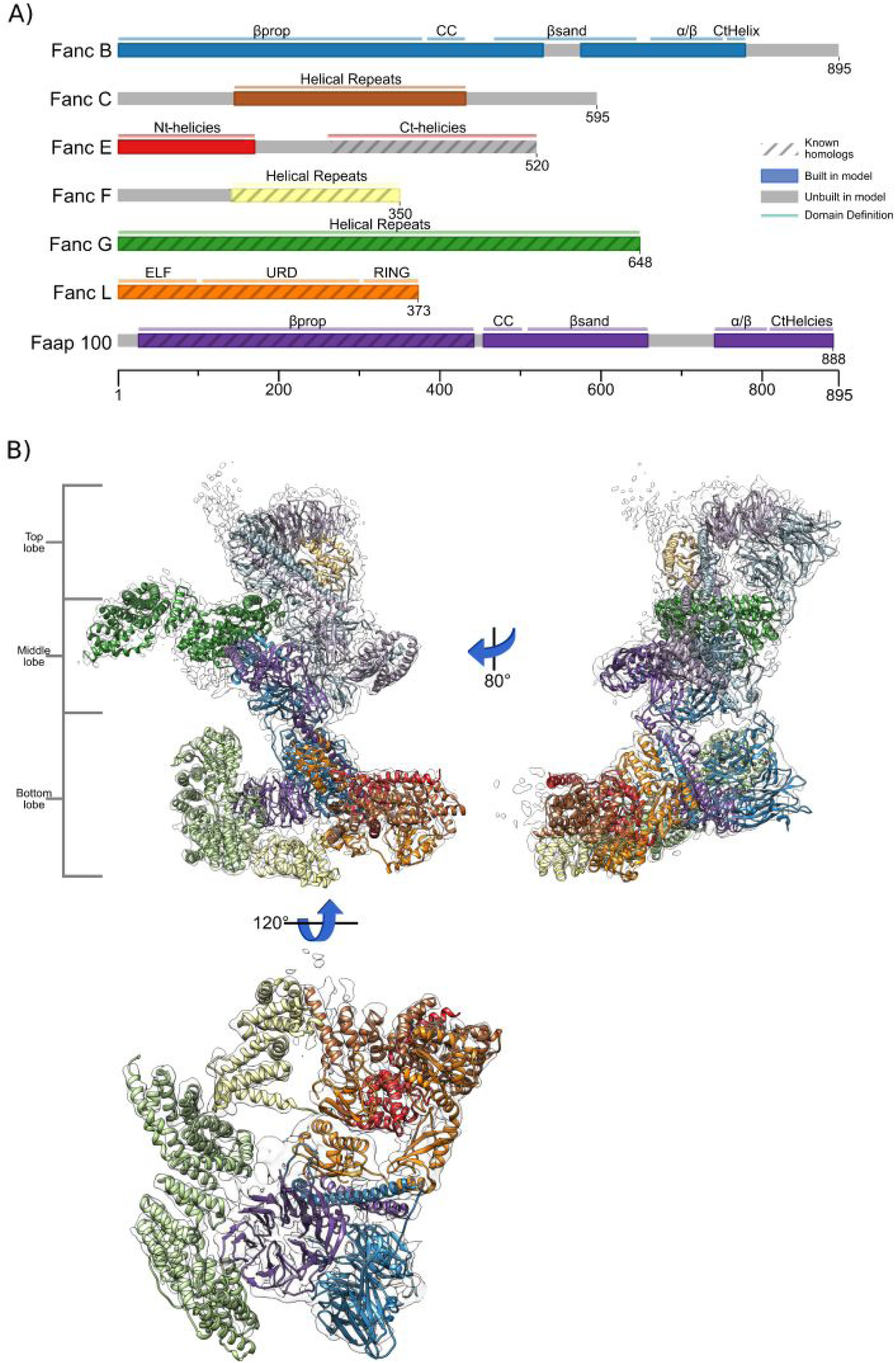
An overview of the FAcc. A) Domain organization of the 7 subunits of FANC. Based on our modeling we find the complex consists of 18 domains, indicated with narrow bars. FANCB and FAAP100 have the same domain organization with a β-propeller(βprop) followed by a long coiled coil (CC), a β-sandwich (βsand), then an α/β domain, followed finally by a C-terminal helical region. FANCC, FANCF, and FANCG are all comprised of a single helical repeat domain, while we find FANCE to have 2 separated helical repeat domains (one N-terminal and one C-terminal.) Finally FANCL is organized as an ELF domain, followed by a URD domain, and then lastly a RING domain. Also indicated is the availability of known structures or homologous proteins throughout the modeling process with striations. Domains with known structures or available homologous proteins used include the C-terminal helices of FANCE, the helical repeats of FANCF, all of FANCG and FANCL, and the β-propeller of FAAP 100. B) Three views of the complete model of the Fanconi Anemia Nuclear Complex as determined by our modeling protocol. Colors are matched to the diagram of subpanel A, with those that have multiple copies (FANCB, FANCG, and FAAP100) having different shades of the subpanel coloring. The orientation of the top, middle, and bottom lobes are indicated.

Using trRosetta-predicted distance distributions, we were able to determine a complete sequence assignment of the full FAcc (Figure 2B), encompassing 5182 residues out of a total expected 6154 residues, or 84% of all sequence, with very little unexplained density (Figure S1). Modelling did not make use of the domain assignments or backbone trace of the prior work. Our model validates much of the putative subunit assignments from the prior study (with minor differences) and provides residue level detail of subunit locations and interactions. The next several sections describe the modelling process, followed by an analysis of our final model.

### Fold trRosetta Models

Our protocol uses multiple sequence alignments (MSAs) for individual proteins as the input to a deep residual-convolutional network which predicts the relative distances and orientations of all residue pairs in the protein. These predictions are applied to a restrained minimization using a Rosetta model building protocol. For FAcc, MSAs were generated for every chain without known homologous structures (homology models were available for portions of FANCE, FANCF, FANCL, and FAAP100, see Figure 2A). Although homologous structures to FANCG also exist, there was significant structural variability within the family, and therefore we modeled it with trRosetta in addition to building homology models.

From the MSAs, domains were manually parsed (see Methods), and models were built using trRosetta (in regions with no known homologs) or comparative modeling (in regions with known homologs). Modelling yielded converged structures for almost all domains (Figure 3A), with typical maximal RMSds over the top models of 2-4Å. Several of the domains that showed poor convergence (two of the domains in FANCB and 2 of the domains in FAAP100) still contained subregions (“converged cores”) with small deviations (2-4 Å) between models; for these cases, unconverged or poorly packed segments of the models were manually trimmed. Three of the domains (the coiled coil domains of FANCB and FAAP100, and the α/β+CTH of FAAP100) were poorly converged with no “converged core”; a modified version of trRosetta (unpublished) in which structural information of distant homologues were used as inputs to the neural network led to well converged models. In total, trRosetta was able to build all 42 attempted domains (Figure S2A) which were used in the following stages of the model building protocol.

**Figure 3.**
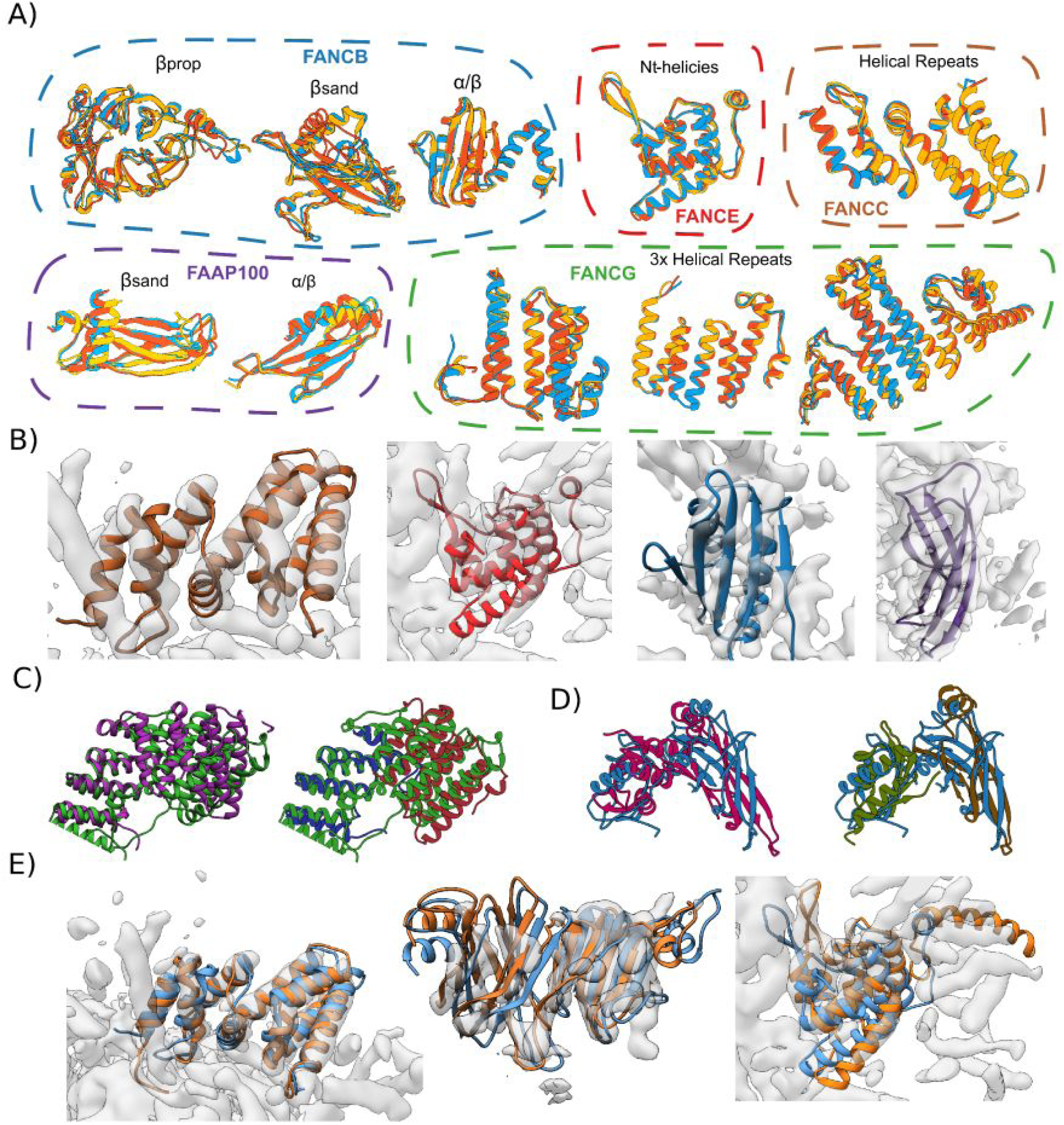
An overview of trRosetta-predicted domains. A)The top three models from trRosetta for ten representative domains indicate tight convergence of modelling. The identity of the domains follows the coloring of Figure 2A. Domains from FANCB, FANCE, FANCC are shown in the top row; those from FAAP 100 and FANCG are shown in the bottom. B) Several examples of trRosetta models docked into density before refinement, showing the role the map plays in validation and selection of models. From left to right (colors match Figure 2A), the helical repeats of FANCC, the N-terminal repeats of FANCE, the α/β domain of FANCB, and the β-sandwich of FAAP100. C-D) Two examples illustrating the importance of domain segmentation when docking trRosetta-generated models. C) The trRosetta model of FANCG (magenta) poorly matches the final structure (green); segmenting this model into two domains (red and blue) shows a much better match, as the individual domain structures are accurate, even though their relative orientation is not. D) Similarly, a trRosetta prediction of the FANCB β-sandwich - α/β domain (pink) is dissimilar from the final structure (blue); splitting into domains (brown and green) shows good overall agreement. E) trRosetta models (blue) generally fit the map well, though some refinement was necessary to maximize agreement with the density (orange).

### Assembling Domains into cryoEM Density

While we found FANCL, FANCF, and FANCE straightforward to manually place into the map, ambiguity for placement of other subunits necessitated a more robust automated assembly procedure. Initially, the top 5 models for each domain were docked using an FFT-accelerated 6D search of the map. A modified version of the Monte Carlo simulated annealing (MC-SA) sampling protocol described in Wang et al. (2015) was then used to identify the non-clashing placement of models that maximizes the overall fit of the complex model to the density. This MC-SA domain assembly assigns a placement or “not found” to each domain, to account for the possibility that either all of our predicted models are incorrect, or that domains are correct but not present in the map. In this way, the map serves not only to orient domains, but as validation for the trRosetta predictions. Some examples of model validation with the map are shown in Figure 3B. Two examples of incorrect predictions (subsequently fixed by splitting models into two domains) are shown in Figure 3C-D. For several domains (the aforementioned coiled coil domains of FANCB and FAAP 100, and the FAAP 100 α/β + CtH domains), manual docking was necessary.

In order to model FAcc in its entirety this Monte Carlo simulated annealing assembly process was applied iteratively: in each round, the converged domains from the previous round were frozen; all unassigned domains were redocked and reassembled. Convergence was assessed by manually inspecting the ten domain assignments with best overall agreement to the density and XL-MS data. Once the iterative process converged (after 5 rounds) -- with the vast majority of the density occupied -- the connections between domains were built and refined in the context of the cryoEM density with RosettaCM (Song et al., 2013). Additionally, placed domains were individually inspected, and poorly placed segments were also rebuilt in RosettaCM.

When refining the final assembled model we found most trRosetta models were quite accurate, often requiring only modest (<6Å RMSd) modifications throughout the refinement process (Figure 2E). Only one placed domain required significant movement: the β-propeller domain of FANCB. To refine this domain, the model was automatically segmented into subdomains (see Methods) and redocked and assembled using the same Monte Carlo procedure before refinement. A comparison between the initial and final structure of the β-propeller after this protocol is shown in Supplemental Video 1.

Finally, for FANCG, a repeat protein for which homologous structures were available, we had additionally used trRosetta for modelling, as predicting changes in repeat geometries can prove challenging. As Figure S3 shows, trRosetta yielded models that contained two long adjacent helices between residues 416 and 491 while homology modeling generated models which contained 4 shorter helices. In assembly, both trRosetta and homology models were considered, and we found that the trRosetta models led to much better agreement between model and map. In contrast, in previous work homology modelling was used for FANCG and we were only able to approximately place ∼280 residues into the map (Shakeel et al., 2019).

### Analysis of final model

With our protocol, we were able to build and assign 5,182 residues (out of 6,557 in the full complex); in previous work only 337 residues were assigned. Still, the protein and domain identities assigned previously were largely consistent with models obtained with this new method: we found similar placements of FANCB, FANCF, FAAP100 and FANCL, as well as one of the two copies of FANCG. While we were unable to identify any density associated with FANCA, trRosetta provided well-converged models (Figure S2A). Combining these models with the crosslinking data, we speculate that a region of unassigned density in the middle of the complex corresponds to FANCA (Figure S4). However, due to the poor quality and incompleteness of the density in this region, we were not able to confidently dock the model into the map.

Our final model reveals that the “bottom lobe” (Figure 2B) contains (using the domain terminology of Figure 2A) FANCB_βprop_, FANCL, FANCE, FANCF, FANCG, and the FAAP100_βprop_. In contrast, in previous work FANCC-FANCE were identified within a region of density that we assign to FANCG. The “middle lobe” of our model consists of 2 copies of FANCB_βsand+α/β+CtH_, a second copy of FANCG, and 2 copies of FAAP100_βsand+α/β+CtH_, all consistent with the previously proposed domain assignment (Shakeel et al., 2019). Finally, the “top lobe” was found to contain a second copy of FANCB_βprop,_ FANCL_ELF_, and a second copy of FAAP100_βprop_. This also is consistent with the hypothesized model of the prior work. Finally, both top and bottom lobes were connected to the middle lobe through a FANCB and FAAP100 intermolecular coiled coil. Thus, in addition to validating much of the domain assignment of previous work, our new model now provides accurate positioning of all protein residues.

### Model Validation

One potential source of model validation arises from the crosslinking data. However, as this data was used in domain assembly, it does not serve as independent validation data. As a measure of confidence, we can still use this data by analyzing the gap between the satisfied crosslinks in our model, and the number satisfied by the *second-best* domain arrangement. In our final model, we see good agreement between crosslinks and model (146/188 total; most of the 834 crosslinks in the full dataset involve FANCA, not present in our model). Of the inter-domain crosslinks, 40/60 (67%) are satisfied to a CA-CA distance of 30 Å, which is regarded as an acceptable distance given the usage of the BS3 crosslinker (Merkley et al., 2014). Freezing the unambiguously placed domains, and redocking the remaining potentially ambiguous domains (see Methods) finds a second-best arrangement only satisfies only 33/63 (52%) of inter-domain crosslinks. This loss in inter-domain crosslink satisfaction provides fairly strong confidence in our final model. Further analysis of the unsatisfied crosslinks reveals that most of the unsatisfied crosslinks (14/19) occur between the C-terminus (residues 103+) of FANCL. Our model suggests that one of the two copies of FANCL in the complex has a disordered C-terminal domain, strongly suggesting that most unsatisfied crosslinks come from this disordered (and possibly dynamic) region.

One particularly strong criteria for model validation is the agreement of the maps to individual domain models The trRosetta models of individual domains were predicted without using density data at all, so rigid-body fitting of these domains into density can be seen as “independent validation.” Aside from domains exhibiting internal symmetry or pseudo-symmetry (FANCB_βprop_, FAAP100_βprop_), we found that trRosetta predictions all matched with real-space correlations of 0.72 or better (FANCC 0.82, FANCE_NtD_ 0.75, FAAP_βsand_ 0.75, FAAP_α/β_ 0.72), while the second-best solution (the best “wrong” solution) has a correlation gap of at least 0.05 in all cases. Subjectively, these second-best, incorrect placements look significantly worse. For our placed domains, this large gap between the best and second-best solutions is quite large, and strongly suggests these domains are unlikely to match this well by random chance.

The overall agreement between the refined model and map is consistent with what we would expect given the resolution of the data. We were able to assess the quality of our model by segmenting it against the three individual focused classification maps (used to generate the composite map used in modeling). We find that the model-map correlation for the bottom and middle reconstructions has an FSC=0.5 crossing at about 7.2Å, while the top reconstruction has an FSC=0.5 crossing of about 7.1Å. The overall model-map FSC curves (Figure S6A) shows that the model-map agreements are worse at higher resolutions for the “bottom” reconstruction than the other two, consistent with local resolution estimates (Figure S6B).

Additionally, we can validate models by mapping human mutation data onto the final structure. Using the Fanconi Anemia Mutation Database (http://www2.rockefeller.edu/fanconi/) database, we identified 30 mutations not identified as benign throughout the complex. While most (22) of these are in the core of protein subunits (likely destabilizing individual subunits), we identify 4 (of the remaining 8) at protein-protein interfaces in our model of the FANCcomplex. Mutations of FANCB 230 and 236 would appear to disturb the interface between the FANCB_βprop_ and FANCG_HR_, while a mutation at FANCB 336 would disturb the interface between the FANCB_βprop_ and the FAAP100_βprop_. Additionally, a mutation to FANCC 295 would likely disturb the interface between FANCC and FANCE. All interface mutations are all marked as magenta spheres in Figure 4A, while non-interface mutations are marked with a tan colored sphere.

**Figure 4.**
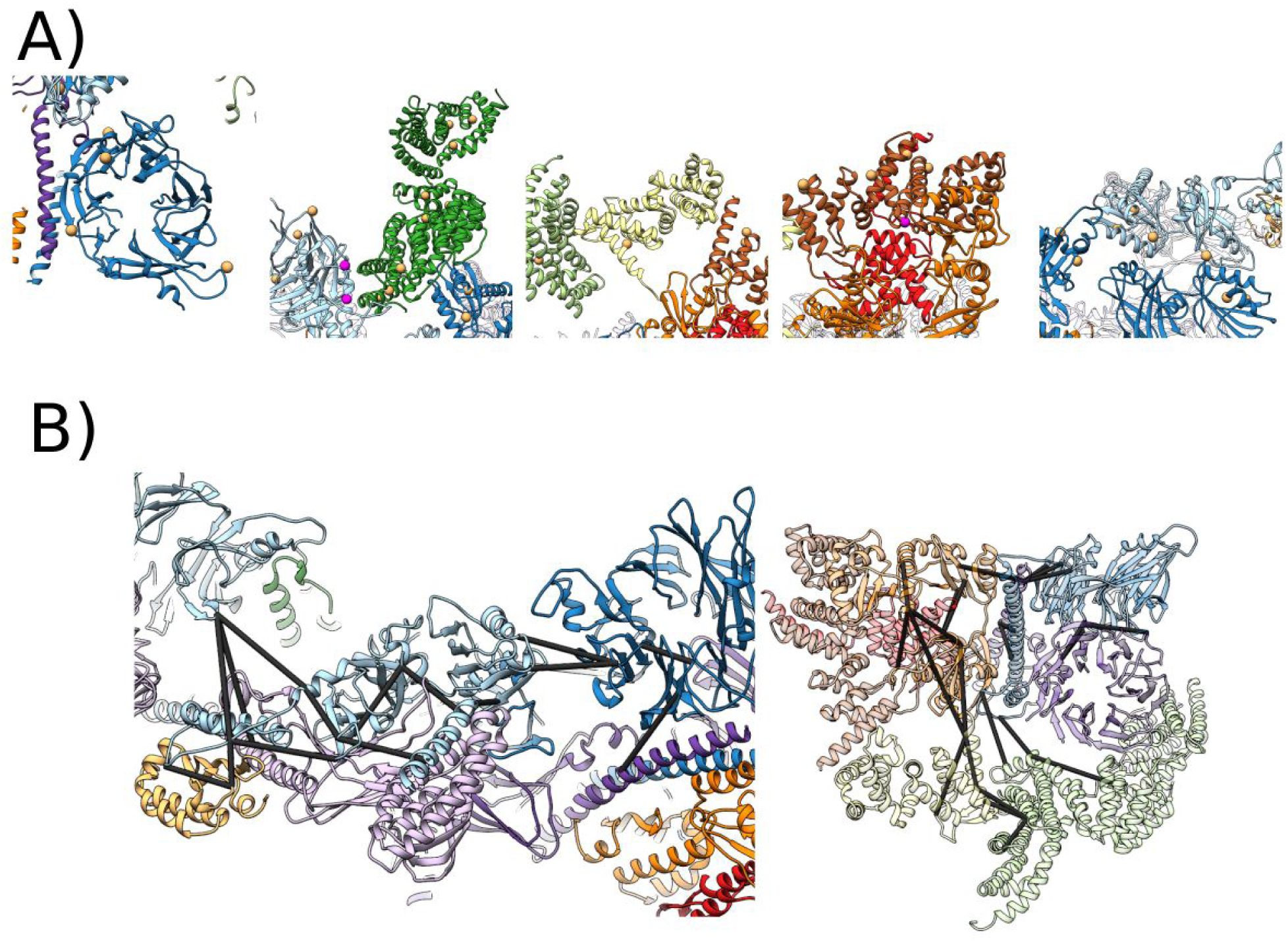
Model validation by mutational and crosslinking data. A) 30 non-benign human mutations mapped to our model of FAcc. All interface mutations are all marked with magenta spheres; non-interface mutations are marked with tan spheres. B) Close up renderings of crosslinks throughout the FAcc model. Black lines indicate crosslinks satisfied (<30 Å) by the final refined structure. Representatives from each crosslink cluster are shown for the middle lobe (left) and the bottom lobe (right).

## Discussion

Here we report a new computational method for determining atomistic models of protein complexes, guided by a subnanometer cryoEM map and cross-linking mass spectrometry data. Using distance distributions predicted from deep residual neural networks, we built accurate models of 42 domains of the FAcc, obviating the necessity for homologous high-resolution structures for interpretation of intermediate resolution maps. This provides a complete picture of the full FAcc, while previous efforts had resulted in atomic models for only three subunits (FANCL, FANCE, and FANCF) in the map. The strong agreement between RosettaTR-predicted models and density (not used in prediction) provides validation of our predictions, as does the model’s consistency with biochemical data, including cross-linking mass spectrometry and mutational studies. Our all-atom model provides molecular insight into the underlying mechanisms of previously reported disease causing mutations, and illustrates the potential of combining intermediate resolution cryoEM density and cutting edge *de-novo* structure prediction.

The challenges faced when determining a model of the FAcc are not unique (Chou et al., 2019; S. J. Kim et al., 2018). As microscopists pursue larger, more difficult, and more dynamic complexes, we will need more computational techniques that are able to build models of subnanometer resolution data with little to no homologous structure information available. While tools have been developed for integrative modelling of structures into subnanometer resolution density, all of these tools require either the existence of homologous structures for domains, or are necessarily low-resolution “domain level” models. Previous attempts to model FAcc resulting in only 387 residues being assigned to the cryoEM data, while the methods described in this paper -- making use of 42 deep-learning guided domain predictions, and a protocol able to infer their arrangement -- were able to increase the number of assigned residues to 5182.

Our approach shows that, while maps at these resolutions are not of sufficient quality to build models by direct chain tracing, the resolution is sufficient enough to assess the tertiary structure and accuracy of predicted models. In the absence of high-resolution homologous structures, the method is able to determine structures to an atomic level of detail. In addition to cryoEM data, we have recently shown that a similar approach can be applied to solve low resolution crystal structure data where traditional molecular replacement techniques were unsuccessful (Bhargava et al., 2020). We expect that the modeling power of trRosetta and related techniques will continue to improve in the future as the number of known sequences increases, coupled with improvements in deep learning methodologies. We anticipate that this combined approach will be an important tool for determining atomic models of protein complexes, particularly when combined with low resolution data sources, enabling accurate protein complex structure determination without the requirement of high resolution data.

## Methods

### Composite map generation

The cryoEM map used for all computation (and displayed in all figures) is a composite map generated from 3 individual focused refinements. The EMDB IDs of these maps are: 10293 (bottom), 10292 (middle), 10291 (top). The maps were combined by first aligning each map to the “bottom” map in UCSF Chimera (Pettersen et al., 2004) using the ‘fit into density’ tool and resampled using “vop resample”. Next, with a custom script the “bottom” map was normalized to density values between 0 and 1, and histogram matching was used to remap the density distribution of the “middle” and “top” map to that of the “bottom.” Finally, a weighted average of the three maps was computed, where the weight of each map’s contribution to the composite map was proportional to the density value in each map at a given point.

Local resolution plots for figure S6B were estimated using ResMap (Kucukelbir et al., 2014) by using the aligned maps generated in the previous step and their respective half maps.

### Subunit multiple sequence alignment generations

In order to model the subunits of FAcc we first generated multiple sequence alignments (MSAs) for every subunit of FAcc with a two-step procedure. In the first stage, four rounds of iterative *HHblits (Steinegger et al., 2019)* (version 3.0.3) searches against the Uniclust30 database (Aug 2018 version) with gradually relaxed e-value cutoffs (10^−80^, 10^−70^, 10^−60^, 10^−40^, 10^−20^) were used to generate an initial alignment. The resulting alignment was then converted to an HMM profile and additional sequences were collected by a single run of *hmmsearch* (version 3.1b2) (Eddy, 1998)against an extensive custom sequence database as described in Wu et al (2020) (Wu et al., 2020); a bit-score threshold of 0.2* (protein length) was used to select significant hits. The composite MSAs were filtered with *hhfilter* at 99% sequence identity and 50% coverage cutoffs.

### trRosetta domain model building

We used trRosetta to predict structures of the following components: FANCA, FANCB, FANCC, FANCE, FANCF, FANCG, FANCL, and Fanconi Anemia core complex Associated Protein (FAAP) 100. trRosetta model building is a two step process where in the initial step a deep residual convolutional neural network is used to generate inter-residue distance and orientation predictions, and in the second step those predictions are used to model a protein of interest (Yang et al., 2020). The MSAs were used as inputs to the neural network which generates residue pair distance distributions in addition to orientation information between all residue pairs. These predictions are then used as input to a custom Rosetta based folding protocol. This protocol works by randomly setting backbone torsions and utilizing random subsets of the predictions as restraints for a centroid (Rosetta’s reduced residue representation) torsional quasi–Newton-based energy minimization (*MinMover*). For each domain 150 centroid models are generated, and then each model is refined with Rostta’s fullatom FastRelax protocol. The results from this refinement are used to sort the models based on the REF2015 score function, and the top 3 models were selected and manually inspected. For all domains except the CC domains and the FAAP FAAP100 α/β + CtH we observed well converged structure, and representative structures from this modeling are shown in figure 3A.

The original trRosetta pipeline was unable to generate converged models for the sequence between the β-propeller regions and the β sandwich regions of FAAP100 and FANCB, and the sequence of FAAP100 α/β + CtH, so we employed a modified version of the network which, in addition to the MSA, also used information on the top 50 putative structural homologs as identified by *HHsearch* against PDB100 database of templates. *HHsearch* hits were converted into 2D network inputs by extracting pairwise distances and orientations from the structure of the template for the matched positions only. Additionally, positional (1D) similarity and confidence scores provided by *HHsearch* as well as backbone torsions were used; we tiled them in both axes of the 2D inputs and stacked with them producing the resulting 2D feature matrix. Features for all unmatched positions were set to zero. Templates were first processed independently by one round of 2D convolutions and then merged together into a single 2D feature matrix using a pixel-wise attention mechanism. This processed feature matrix was then concatenated with the features extracted from the MSA as in the original trRosetta network; the architecture of the upstream part of the network remained unchanged. For the CC domains this improved the quality of models for the β-propellers as well as models for the extended helices C-terminal to the β-propellers. For FAAP100 α/β + CtH we modeled FAAP100 CC+βsand+α/β+CtH with this modified version and found strong convergence for all of the domains. The coiled coil domains of FANCB and FAAP, and the FAAP100 α/β and CtH were manually extracted for use in the next stages. The results from this modified version of trRosetta are shown in figure S2B.

### Inferring domain boundaries

To infer domain boundaries we used the MSAs as initial guidelines by adding cutpoints at residues with poor alignment coverage. Using these domain definitions we then performed structural modeling of the domains and used these models to manually split the sequence further based on the observed convergence, and trimmed away floppy regions. The following domain boundaries were determined:

- FANCA: [71-260], [251-500], [500-651], [1011-1210]
- FANCB: [1-235], [1-370], [231-365], [441-660], [441-780], [466-626], [651-770], [665-755]
- FANCC : [1-175], [1-335], [176-335], [331-570]
- FANCE: [1-150], [261-520]
- FANCF: [1-130], [121-350], [142-350]
- FANCG: [1-175], [1-320], [181-320], [201-435], [204-315], [321-648], [350-564]
- FANCL: [1-100], [2-91], [101-205], [101-300], [104-373], [191-300], [301-373]
- FAAP100: [1-200], [1-300], [28-442], [186-480], [301-480], [491-615], [491-820], [510-609], [711-820], [717-820]

Then, based on the availability of homologous structures in these regions either RosettaCM (Song et al., 2013) (if homologous structures were available) or trRosetta (Yang et al., 2020) (if homologous structures were not available) were used to generate models for each domain.

### RosettaCM domain model building

We modeled FANCE_CtH_, FANCF_hr_, FANCG, FANCL_ELF+URD+RING_ hr, and FAAP100_βprop_ usingRosettaCM (Song et al., 2013). The following templates were used for each subunit:

- FANCE: 2ilr (chain A)
- FANCF_HR_: 2iqc (chain A)
- FANCG: 6eou (chain A), 2xpi (chain A), 3hym (chain J), 3cvp (chain A), 4rg9 (chain A), 5dse (chain A), 3fp2 (chain A), 5orq (chain A), 5i9f (chain A), 4g1t (chain B), 3ieg (chain A), 2y4t (chain A), 5aio (chain A), 4pjr (chain A), 1fch (chain B), 4zlh (chain B), 2gw1 (chain A), 6c9m (chain C), 3u4t (chain A), 4buj (chain B)
- FANCL_ELF+URD+RING_: 3k1l (chain B), 4zdt (chain A), 4ccg (chain Y), 1vyx (chain A), 5o6c (chain A), 2d8s (chain A)
  - (the resulting models were segmented more before docking)
- FAAP100_βprop_: 4ggc (chain A), 5opt (chain p), 5xyi (chain g), 6chg (chain A), 5oql (chain F), 2pbi (chain D), 1r5m (chain A), 6eoj (chain D), 6f9n (chain B), 5m89 (chain B), 3odt (chain B), 5a31 (chain R), 5m23 (chain A), 5kdo (chain B)

For each of the above 200 models were generated, using the command line:

~~~
   rosetta_scripts \
   -in:file:fasta FANCX.fasta \
   -parser:protocol hybridize.xml \
   -relax:jump_move true \
   -default_max_cycles 200 \
   -beta_cart \
   -relax:dualspace
~~~

The input XML file (hybridize.xml) is shown below:

~~~
   <ROSETTASCRIPTS>
     <SCOREFXNS>
      <ScoreFunction name=“stage1” weights=“score3”>
         <Reweight scoretype=“atom_pair_constraint” weight=“0.1”/>
      </ScoreFunction>
      <ScoreFunction name=“stage2” weights=“score4_smooth_cart”>
         <Reweight scoretype=“atom_pair_constraint” weight=“0.1”/>
       </ScoreFunction>
     <ScoreFunction name=“fullatom” weights=“beta_cart”>
       <Reweight scoretype=“atom_pair_constraint” weight=“0.1”/>
     </ScoreFunction>
   </SCOREFXNS>
   <MOVERS>
     <Hybridize name=“hybridize” stage1_scorefxn=“stage1”
         stage2_scorefxn=“stage2” fa_scorefxn=“fullatom” batch=“1” >
       <Template pdb=“000_2ilr_A.pdb” cst_file=“AUTO” weight=“1.0” />
     </Hybridize>
   </MOVERS>
   <PROTOCOLS>
      <Add mover=“hybridize” />
   </PROTOCOLS>
   <OUTPUT scorefxn=“fullatom” />
  </ROSETTASCRIPTS>
~~~

### Docking of domain models into density

A new Rosetta tool, *`dock_pdb_into_density`*, and a wrapper script (dgdp,py, for density-guided domain placement) were used for the initial assembly of models. Briefly, dock_pdb_into_density uses a FFT-accelerated six-dimensional search to find the rigid body placements of a molecule that maximize overlap between model and map. For all domains that were modelled, we began by docking the top 3 models into density using dock_pdb_into_density. This method carries out FFT convolutions in rotational space, explicitly enumerating over translations, the method identified 50,000 points with high density (and >2 Å apart). For each domain, all solutions were combined, and the top 1000 were filtered, and rigid-body minimized in Rosetta using a masked correlation function (DiMaio et al., 2009). After minimization, results were filtered for redundancy (using an 11 Å RMS cutoff) and the top 200 solutions were selected.

The following command line carries out the procedure for FANCC:

~~~
python dgdp.py \
    --executable ∼/Rosetta/dgdp/source/bin/dgdp.static.linuxgccrelease \
    --point_radius 3.0 \
    --n_output 200 \
    --n_filtered 1000 \
    --n_to_search 50000 \
    --mapfile fancc_map.mrc \
    --clust_radius 11 \
    --mapreso 5 \
    --pdbs input_fancc_* .pdb \
    --name fancc \
    --cores 10 \
    --min \
    --constrain_refinement 10 \
    --queue medium \
    --final_result_name FANC_C0
~~~

. Typical cpu usage for docking one model is highly dependent on the size of the density map and the number of residues in the model, but for FAcc we generally see docking take 5.5 hours total cpu time. However this is highly parallelizable, and by using the python *dask* library (Dask Development Team, 2016) with ample computing resources the total time taken can be reduced significantly (54 minutes with 10 CPUs).

### Docked domain assembly

Given the docked domains of the previous section, we use a modified version of the Monte Carlo simulated annealing (MC-SA) sampling protocol described in Wang et al. (2015) to build a model of the complex. Briefly, the protocol uses the top 200 placements for each model from our docking protocol, in addition to the crosslinking data, in order to determine a set of domain placements most consistent with all available data. This MC-SA domain assembly assigns a placement or “not found” to each domain, to account for the possibility that either all of our predicted models are incorrect, or that domains are correct but not present in the map. Consistency is measured through the function (where *d*_*N*_ is all domains):

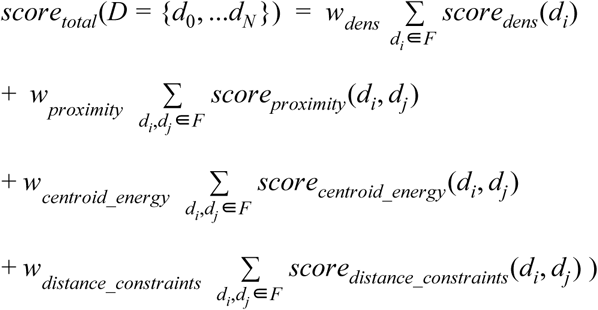

Where *score*_*dens*_ measures the fit of the selected domains to the density and the other terms assess interactions between all domains. The term *score*_*proximity*_ validates that when two domains are part of the same peptide chain and not overlapping they are placed within a distance that is closable by a later built peptide linker. The *score*_*centroid energy*_ term is Rosetta’s centroid energy score term which is a coarse grained representation that is used to verify the quality of domain-domain interfaces, as well as screen for clashing placements. The centroid energy between two domains is evaluated by using Rosetta to combine the two domains into a single system (Pose), evaluate the system’s energy, then spatially separating the two domains and again evaluating the energy of the system. The former is subtracted by the latter, and this is used as the unweighted *centroid_energy* score. Finally the *score*_*distance constraint*_ term serves as a way to incorporate experimental data such as cross linking mass-spectrometry data, and assesses the satisfaction of these constraints. The inter-domain geometry terms are assessed as follows:

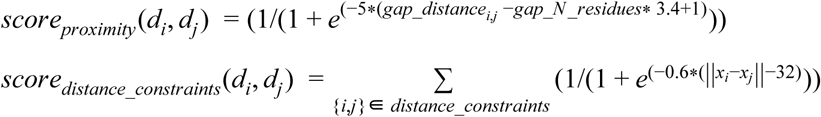

Weights were determined by fitting on a training set with synthetic 10 Å cryoEM data, and the weights used were *w*_*dens*_ = 260, *w*_*distance_constraint*_ = 30,000, *w*_*centroid_energy*_=150, *w*_*proximity*_= 1,000.

Using the scoring function above, we evaluate the consistency of the results from docking for all domain-domain pairs. Prior to scorefunction evaluation a custom pairwise interface optimization protocol is applied: domains are slid along an axis through each domains’ center of mass to be in contact, but not clashing with each other, moving no more than 5 Å. If after this, domains are still clashing (defined as Rosetta vdw score > 1500), we remove all clashing residues (Rosetta vdw score > 3) with: a) no secondary structure, and b) surface exposure (less than 10 Cα’s within 12Å), and rescored. This is then followed by breaking both domains into subdomains (using a reimplementation of DDomainParse (Zhou et al., 2007) in Rosetta), and rigid-body minimizing these domains with respect to the energy function above.

Once all pairs of domains have been refined, and their refined inter-domain energies have been computed, Monte Carlo simulated annealing (MC-SA) sampling is carried out. Each MC-SA move reassigns one domain to either another placement, or “no placement.” We carry out 200,000 steps of MC-SA sampling, ending at a temperature of kT=1. 50,000 independent trajectories are carried out and the top ten scoring assignments are assessed for convergence. Convergence is assessed by manually inspecting the ten domain assignments for domains that were present in a majority of the models. This process is applied iteratively; after each round of assembly, domains that converged in location are locked into place, where convergence was assessed, the density occupied by converged domains was removed from the density map, and unassigned domains were re-docked and used as inputs for the next round.

In the case of FAcc, the iterative process progressed as follows: After the first round FANCB_α/β_, FANCC, FANCE_NtD_, FANCF, 6 helices of FANCG (204-315, and FANCL were found to converged and were locked into place. After the second round, 8 helices of FANCG (350-564) were locked into place. After this round, the density associated with the two β-propellers was segmented out, and both the FANCB and FAAP β-propellers were docked into this segmented density, which were used as inputs for the next round. During the third round, the β-propellers of FANCB and FAAP100 and the two coiled coils of FANCB and FAAP100 both were frozen into place. In the fourth round, we found converged placement of FANCB_βsand_. After this round utilizing the remaining models and density, we manually docked the FAAP 100_Bsand_, FAAP 100_α/β_, and FAAP 100_CtH_ domains into the density.

### Structure Finalization

In order to finalize the structure, all of the placed domains from the previous step were combined and linked together with RosettaCM (Song et al., 2013). There were five areas that required directed rebuilding with Rosetta, the FANCB β-propeller, the FANCB loop between the CC helix and β-sandwich, The FANCB C-terminal helix, FANCG, and the FAAP 100_α/β+CtH_ domain.

The FANCB β-propeller as solved by trRosetta was very similar to the density but the spacing between each propeller blade was off by a significant enough margin to make it difficult for RosettaCM to properly minimize into the density. Therefore, we ran an automatic domain splitting script, and used *dock_pdb_into_density* to place subdomains of the propeller into the β-propeller density, assembled using the protocol described above.

The FANCB helix to β sandwich loop required excessive sampling to build due to its length (41 residues) and lack of density. The density around this area was segmented, and iterative hybridize (H. Park et al., 2018) was run with the initial amount of structures generated being 5000 followed by 4 rounds each generating 100 structures.

The FANCB extended helix built with trRosetta was added, using Chimera’s fit into map tool, after the α/β domain of FANCB had been placed. This was done because of the unambiguous density leading from the α/β extending to helical density which made.

For FANCG, assembly placed only two domains, corresponding to residues 204-315 and 350-564. The remaining structure was built in the following way. First, The N terminal domain (1-204) was well converged in trRosetta, and was manually docked into the map by aligning to overlapping residues in one of the placed domains (residues 200-230 overlapped between the two). The same process was carried out with the C-terminal domain (residues 565-648), where the overlapping residues used were 551-562. These placements were validated by manually inspecting fit to density.

The FAAP100_α/β_ domain posed a particularly difficult problem due to low local resolution and poor connectivity (this domain is preceded by a long unstructured loop). Due to these ambiguities, FAAP100_CtH_ models had to be manually aligned to the density (using Chimera’s fit into map). Full-length trRosetta models were used as a reference for placement.

Finally, after refining the structure with RosettaCM, we applied fragment based structural refinement (Wang et al., 2016), and selected the top scoring model as our final model.

## Data Accessibility

All methods are available in Rosetta releases after 2020.12.

The model is deposited in pdb-dev with accession id XXX.

The crosslinking data that was as distance constraints during this process is located at the PRIDE database with accession code PXD014282.

---

**Figure S1.**
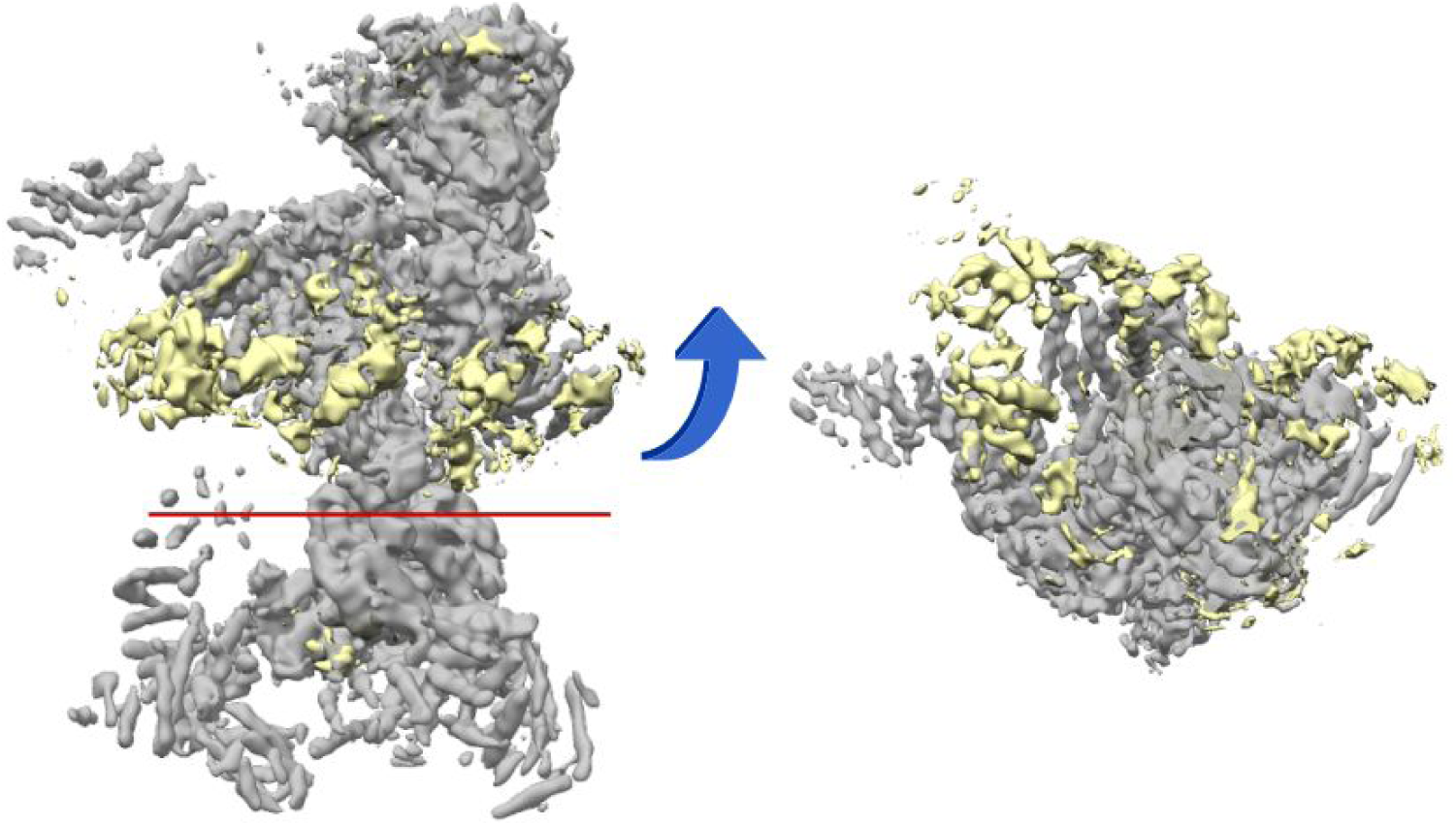
An overlay of the density map used to build the FAcc complex (Grey), with the density not within 3.5 Å of the model highlighted in yellow. We were unable to build into the yellow density due to the low resolution, and lack of continuity between regions.

**Figure S2.**
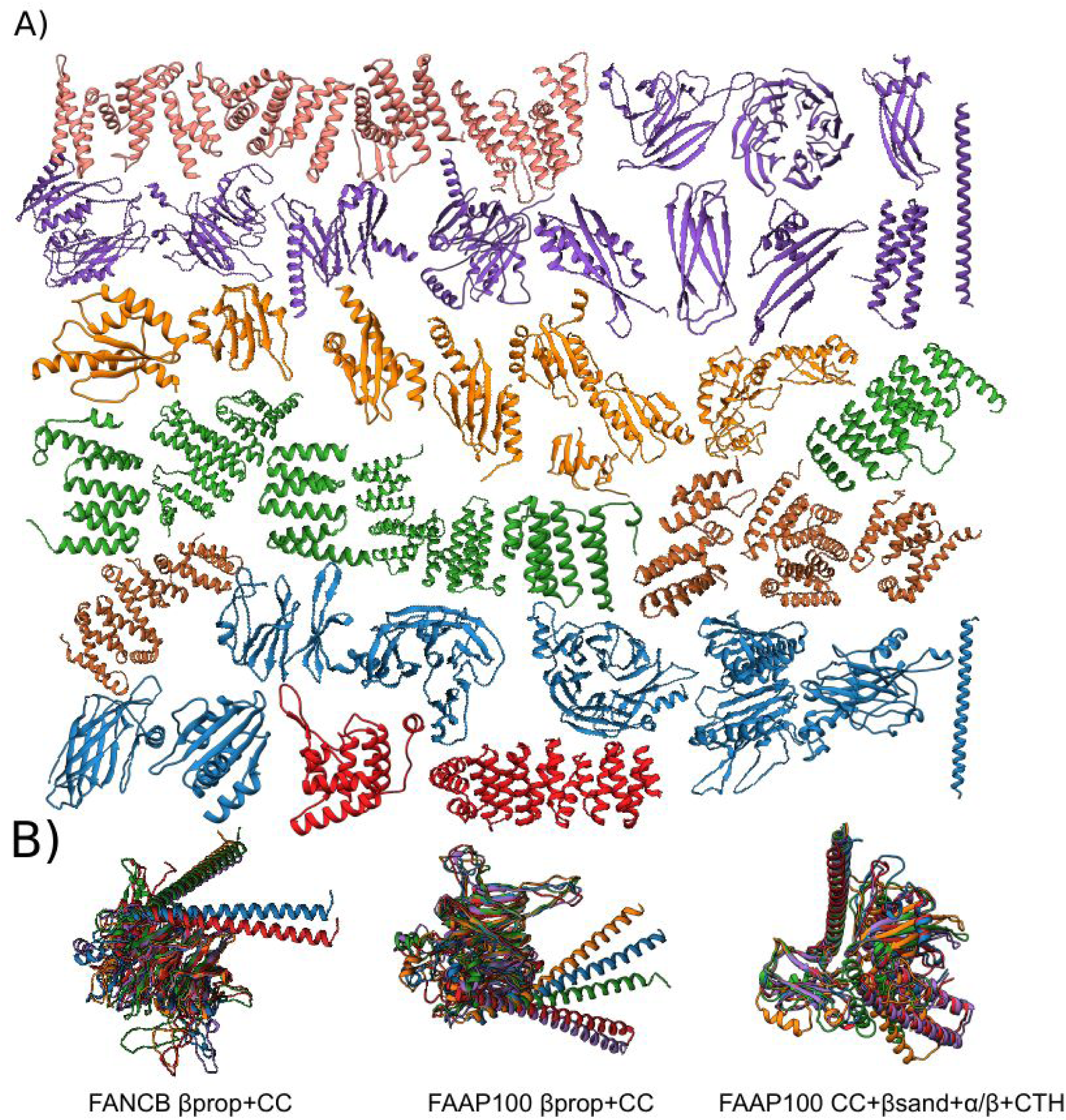
Top scoring models for all of the domains built in order to build the FAcc. A) All domains that were individually docked into the density. Colors match Figure 2A. B) All models built by the novel implementation of trRosetta that uses distant homology information in order to improve prediction accuracy. In these three cases, the original implementation of trRosetta was unable to generate well converged models for the combinations shown.

**Figure S3.**
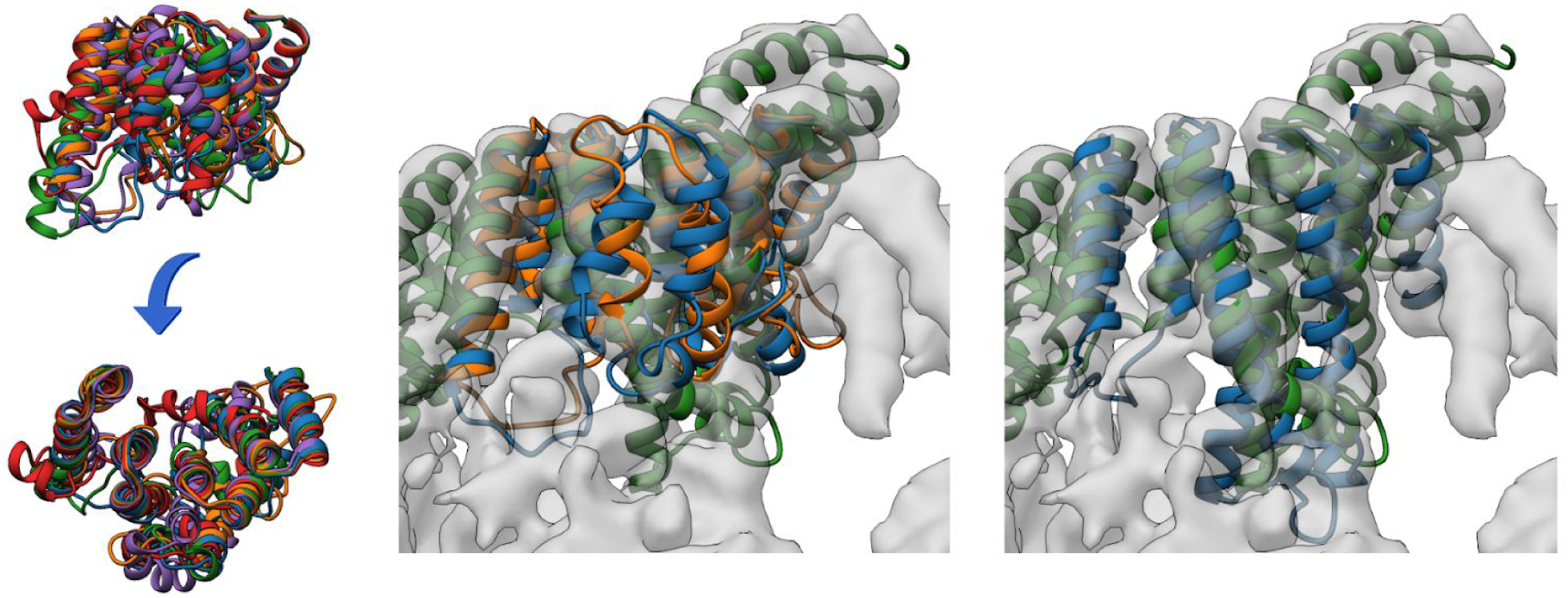
Top 5 scoring ensemble from RosettaCM from FANCG (left), with the top 2 models overlaid on the final structure of FANCG (middle). The right shows the model made by trRosetta, which fits the density significantly better than the RosettaCM models

**Figure S4.**
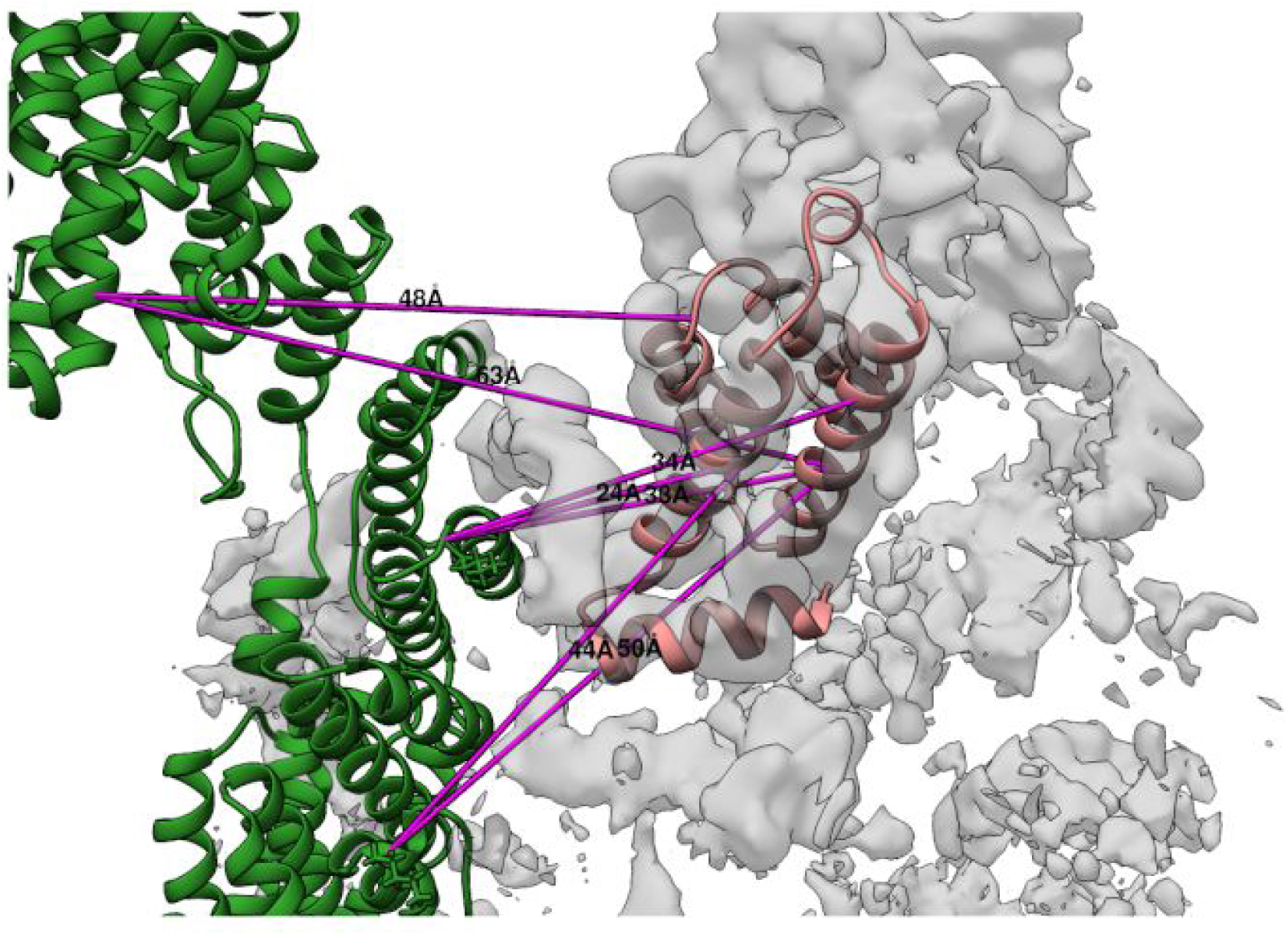
A speculative location for the N-terminal domain of FANCA. We hypothesize, based on the trRosetta models fit to the sparse electron density, and crosslink satisfaction that the N-terminal domain of FANCA rests at or near this location.

**Figure S5.**
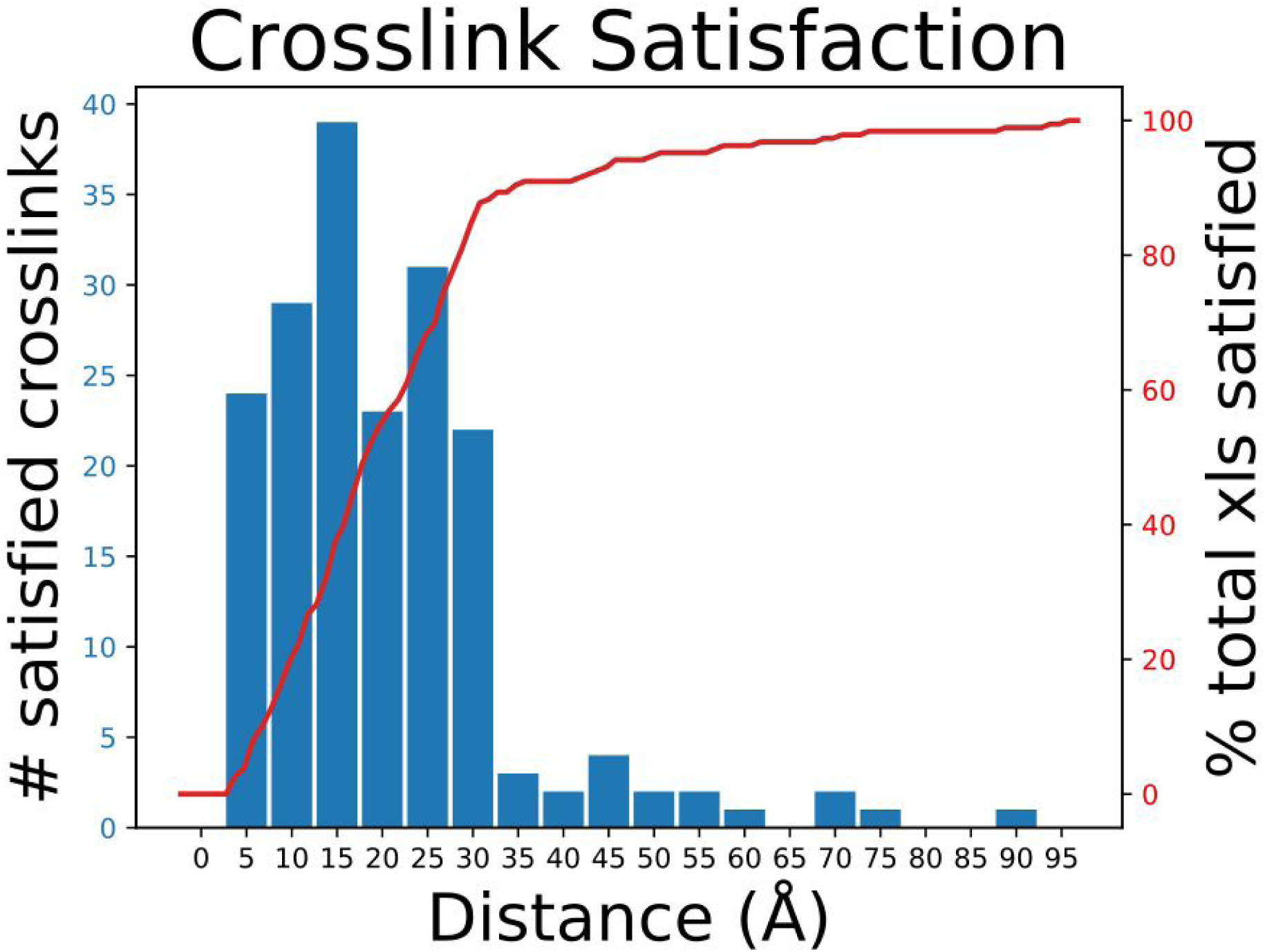
Evaluation of crosslink satisfaction across our final model. We see approximately 77% satisfaction of all crosslinks at a cutoff of 30Å which is acceptable given our BS3 crosslinker. The histogram in blue plots the number of crosslinks satisfied in 5 Å bins (0-5, 5-10, …) and is plotted against the left Y axis. The line plot in red is a continuous measure of the percent of the total number of crosslinks satisfied in our model (right Y axis) as a function of their evaluated distance in Å.

**Figure S6.**
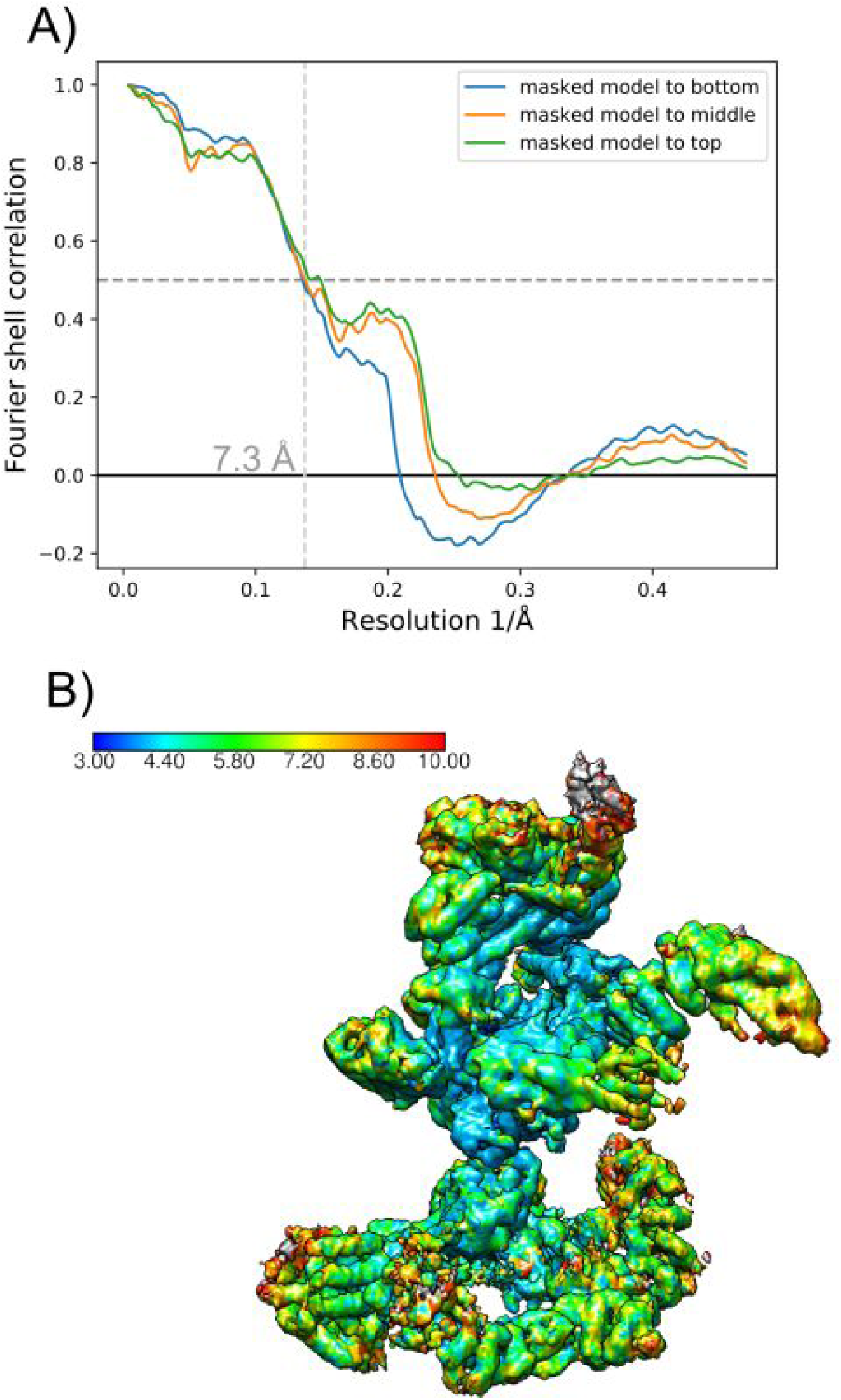
Evaluation of model from map. A) FSC plot of our model segmented to the three maps that generated the composite map used throughout modeling. B) Local resolution plot of the map used during modeling.

## Supplemental video 1

Supp_video_01_fanc.mp4

## Notes

### Competing Interest Statement

The authors have declared no competing interest.

## References

Bhargava, H. K., Tabata, K., Byck, J. K., Hamasaki, M., Farrell, D. P., Anishchenko, I., DiMaio, F., Im, Y.-J., Yoshimori, T., & Hurley, J. H. (2020). *Structural basis for autophagy* inhibition by the human Rubicon-Rab7 complex [Preprint]. Cell Biology. https://doi.org/10.1101/2020.04.18.048462

Bonomi, M., Hanot, S., Greenberg, C. H., Sali, A., Nilges, M., Vendruscolo, M., & Pellarin, R. (2019). Bayesian Weighing of Electron Cryo-Microscopy Data for Integrative Structural Modeling. Structure, 27 (1), 175-188.e6. https://doi.org/10.1016/j.str.2018.09.011

Chen, M., Baldwin, P. R., Ludtke, S. J., & Baker, M. L. (2016). De Novo modeling in cryo-EM density maps with Pathwalking. Journal of Structural Biology, 196 (3), 289–298. https://doi.org/10.1016/j.jsb.2016.06.004

Cheng, L., Zhu, J., Hui, W. H., Zhang, X., Honig, B., Fang, Q., & Zhou, Z. H. (2010). Backbone Model of an Aquareovirus Virion by Cryo-Electron Microscopy and Bioinformatics. Journal of Molecular Biology, 397 (3), 852–863. https://doi.org/10.1016/j.jmb.2009.12.027

Cheng, Y., & Walz, T. (2009). The Advent of Near-Atomic Resolution in Single-Particle Electron Microscopy. Annual Review of Biochemistry, 78 (1), 723–742. https://doi.org/10.1146/annurev.biochem.78.070507.140543

Dask Development Team. (2016). Dask: Library for dynamic task scheduling URL https://dask.org. https://dask.org

DiMaio, F., Tyka, M. D., Baker, M. L., Chiu, W., & Baker, D. (2009). Refinement of Protein Structures into Low-Resolution Density Maps Using Rosetta. Journal of Molecular Biology, 392 (1), 181–190. https://doi.org/10.1016/j.jmb.2009.07.008

Eddy, S. R. (1998). Profile hidden Markov models. Bioinformatics, 14 (9), 755–763. https://doi.org/10.1093/bioinformatics/14.9.755

Gatsogiannis, C., Lang, A. E., Meusch, D., Pfaumann, V., Hofnagel, O., Benz, R., Aktories, K., & Raunser, S. (2013). A syringe-like injection mechanism in Photorhabdus luminescens toxins. Nature, 495 (7442), 520–523. https://doi.org/10.1038/nature11987

Hryc, C. F., Chen, D.-H., & Chiu, W. (2011). Near-atomic resolution cryo-EM for molecular virology. Current Opinion in Virology, 1 (2), 110–117. https://doi.org/10.1016/j.coviro.2011.05.019

Janssen, M. E. W., Takagi, Y., Parent, K. N., Cardone, G., Nibert, M. L., & Baker, T. S. (2015). Three-Dimensional Structure of a Protozoal Double-Stranded RNA Virus That Infects the Enteric Pathogen Giardia lamblia. Journal of Virology, 89 (2), 1182–1194. https://doi.org/10.1128/JVI.02745-14

Kandathil, S. M., Greener, J. G., & Jones, D. T. (2019). Prediction of interresidue contacts with DeepMetaPSICOV in CASP13. Proteins: Structure, Function, and Bioinformatics, 87 (12), 1092–1099. https://doi.org/10.1002/prot.25779

Kim, D. E., DiMaio, F., Yu-Ruei Wang, R., Song, Y., & Baker, D. (2014). One contact for every twelve residues allows robust and accurate topology-level protein structure modeling: Contact Guided Protein Structure Prediction. Proteins: Structure, Function, and Bioinformatics, 82, 208–218. https://doi.org/10.1002/prot.24374

Klink, B. U., Gatsogiannis, C., Hofnagel, O., Wittinghofer, A., & Raunser, S. (2020). Structure of the human BBSome core complex. ELife, 9. https://doi.org/10.7554/eLife.53910

Kovacs, J. A., Galkin, V. E., & Wriggers, W. (2018). Accurate flexible refinement of atomic models against medium-resolution cryo-EM maps using damped dynamics. BMC Structural Biology, 18 (1), 12. https://doi.org/10.1186/s12900-018-0089-0

Kube, S., Kapitein, N., Zimniak, T., Herzog, F., Mogk, A., & Wendler, P. (2014). Structure of the VipA/B Type VI Secretion Complex Suggests a Contraction-State-Specific Recycling Mechanism. Cell Reports, 8 (1), 20–30. https://doi.org/10.1016/j.celrep.2014.05.034

Kucukelbir, A., Sigworth, F. J., & Tagare, H. D. (2014). Quantifying the local resolution of cryo-EM density maps. Nature Methods, 11 (1), 63–65. https://doi.org/10.1038/nmeth.2727

Lyumkis, D. (2019). Challenges and opportunities in cryo-EM single-particle analysis. Journal of Biological Chemistry, 294 (13), 5181–5197. https://doi.org/10.1074/jbc.REV118.005602

Merkley, E. D., Rysavy, S., Kahraman, A., Hafen, R. P., Daggett, V., & Adkins, J. N. (2014). Distance restraints from crosslinking mass spectrometry: Mining a molecular dynamics simulation database to evaluate lysine–lysine distances. Protein Science : A Publication of the Protein Society, 23 (6), 747–759. https://doi.org/10.1002/pro.2458

Nugent, T., & Jones, D. T. (2012). Accurate de novo structure prediction of large transmembrane protein domains using fragment-assembly and correlated mutation analysis. Proceedings of the National Academy of Sciences, 109 (24), E1540–E1547. https://doi.org/10.1073/pnas.1120036109

Ovchinnikov, S., Kamisetty, H., & Baker, D. (2014). Robust and accurate prediction of residue–residue interactions across protein interfaces using evolutionary information. ELife, 3. https://doi.org/10.7554/eLife.02030

Park, H., Ovchinnikov, S., Kim, D. E., DiMaio, F., & Baker, D. (2018). Protein homology model refinement by large-scale energy optimization. Proceedings of the National Academy of Sciences, 115 (12), 3054–3059. https://doi.org/10.1073/pnas.1719115115

Park, Y.-J., Lacourse, K. D., Cambillau, C., DiMaio, F., Mougous, J. D., & Veesler, D. (2018). Structure of the type VI secretion system TssK–TssF–TssG baseplate subcomplex revealed by cryo-electron microscopy. Nature Communications, 9 (1). https://doi.org/10.1038/s41467-018-07796-5

Pettersen, E. F., Goddard, T. D., Huang, C. C., Couch, G. S., Greenblatt, D. M., Meng, E. C., & Ferrin, T. E. (2004). UCSF Chimera—A visualization system for exploratory research and analysis. Journal of Computational Chemistry, 25 (13), 1605–1612. https://doi.org/10.1002/jcc.20084

Schoebel, S., Mi, W., Stein, A., Ovchinnikov, S., Pavlovicz, R., DiMaio, F., Baker, D., Chambers, M. G., Su, H., Li, D., Rapoport, T. A., & Liao, M. (2017). Cryo-EM structure of the protein-conducting ERAD channel Hrd1 in complex with Hrd3. Nature, 548 (7667), 352–355. https://doi.org/10.1038/nature23314

Segura, J., Sanchez-Garcia, R., Tabas-Madrid, D., Cuenca-Alba, J., Sorzano, C. O. S., & Carazo, J. M. (2016). 3DIANA: 3D Domain Interaction Analysis: A Toolbox for Quaternary Structure Modeling. Biophysical Journal, 110 (4), 766–775. https://doi.org/10.1016/j.bpj.2015.11.3519

Senior, A. W., Evans, R., Jumper, J., Kirkpatrick, J., Sifre, L., Green, T., Qin, C., Žídek, A., Nelson, A. W. R., Bridgland, A., Penedones, H., Petersen, S., Simonyan, K., Crossan, S., Kohli, P., Jones, D. T., Silver, D., Kavukcuoglu, K., & Hassabis, D. (2020). Improved protein structure prediction using potentials from deep learning. Nature, 577 (7792), 706–710. https://doi.org/10.1038/s41586-019-1923-7

Shakeel, S., Rajendra, E., Alcón, P., O’Reilly, F., Chorev, D. S., Maslen, S., Degliesposti, G., Russo, C. J., He, S., Hill, C. H., Skehel, J. M., Scheres, S. H. W., Patel, K. J., Rappsilber, J., Robinson, C. V., & Passmore, L. A. (2019). Structure of the Fanconi anaemia monoubiquitin ligase complex. Nature, 575 (7781), 234–237. https://doi.org/10.1038/s41586-019-1703-4

Snijder, J., Schuller, J. M., Wiegard, A., Lössl, P., Schmelling, N., Axmann, I. M., Plitzko, J. M., Förster, F., & Heck, A. J. R. (2017). Structures of the cyanobacterial circadian oscillator frozen in a fully assembled state. Science, 355 (6330), 1181–1184. https://doi.org/10.1126/science.aag3218

Song, Y., DiMaio, F., Wang, R. Y.-R., Kim, D., Miles, C., Brunette, T., Thompson, J., & Baker, D. (2013). High-Resolution Comparative Modeling with RosettaCM. Structure, 21 (10), 1735–1742. https://doi.org/10.1016/j.str.2013.08.005

Steinegger, M., Meier, M., Mirdita, M., Vöhringer, H., Haunsberger, S. J., & Söding, J. (2019). HH-suite3 for fast remote homology detection and deep protein annotation. BMC Bioinformatics, 20 (1). https://doi.org/10.1186/s12859-019-3019-7

Stuttfeld, E., Aylett, C. H., Imseng, S., Boehringer, D., Scaiola, A., Sauer, E., Hall, M. N., Maier, T., & Ban, N. (2018). Architecture of the human mTORC2 core complex. ELife, 7. https://doi.org/10.7554/eLife.33101

Terashi, G., & Kihara, D. (2018). De novo main-chain modeling for EM maps using MAINMAST. Nature Communications, 9 (1). https://doi.org/10.1038/s41467-018-04053-7

Terwilliger, T. C., Adams, P. D., Afonine, P. V., & Sobolev, O. V. (2018). A fully automatic method yielding initial models from high-resolution cryo-electron microscopy maps. Nature Methods, 15 (11), 905–908. https://doi.org/10.1038/s41592-018-0173-1

van Zundert, G. C. P., Melquiond, A. S. J., & Bonvin, A. M. J. J. (2015). Integrative Modeling of Biomolecular Complexes: HADDOCKing with Cryo-Electron Microscopy Data. Structure, 23 (5), 949–960. https://doi.org/10.1016/j.str.2015.03.014

Wang, R. Y.-R., Song, Y., Barad, B. A., Cheng, Y., Fraser, J. S., & DiMaio, F. (2016). Automated structure refinement of macromolecular assemblies from cryo-EM maps using Rosetta. ELife, 5. https://doi.org/10.7554/eLife.17219

Webb, B., Viswanath, S., Bonomi, M., Pellarin, R., Greenberg, C. H., Saltzberg, D., & Sali, A. (2018). Integrative structure modeling with the Integrative Modeling Platform: Integrative Structure Modeling with IMP. Protein Science, 27 (1), 245–258. https://doi.org/10.1002/pro.3311

Wu, Q., Peng, Z., Anishchenko, I., Cong, Q., Baker, D., & Yang, J. (2020). Protein contact prediction using metagenome sequence data and residual neural networks. Bioinformatics, 36 (1), 41–48. https://doi.org/10.1093/bioinformatics/btz477

Xu, J. (2019). Distance-based protein folding powered by deep learning. Proceedings of the National Academy of Sciences, 116 (34), 16856–16865. https://doi.org/10.1073/pnas.1821309116

Yang, J., Anishchenko, I., Park, H., Peng, Z., Ovchinnikov, S., & Baker, D. (2020). Improved protein structure prediction using predicted interresidue orientations. Proceedings of the National Academy of Sciences, 117 (3), 1496–1503. https://doi.org/10.1073/pnas.1914677117

Zheng, W., Li, Y., Zhang, C., Pearce, R., Mortuza, S. M., & Zhang, Y. (2019). Deep-learning contact-map guided protein structure prediction in CASP13. Proteins: Structure, Function, and Bioinformatics, 87 (12), 1149–1164. https://doi.org/10.1002/prot.25792

Zhou, H., Xue, B., & Zhou, Y. (2007). DDOMAIN: Dividing structures into domains using a normalized domain–domain interaction profile. Protein Science : A Publication of the Protein Society, 16 (5), 947–955. https://doi.org/10.1110/ps.062597307

